# New insights from short and long reads sequencing to explore cytochrome *b* variants of *Plasmopara viticola* populations collected in vineyard and related to resistance to complex III inhibitors

**DOI:** 10.1101/2022.04.29.490075

**Authors:** Semcheddine Cherrad, Benjamin Gillet, Julien Dellinger, Lalie Bellaton, Pascale Roux, Catalina Hernandez, Hervé Steva, Lauriane Perrier, Sandrine Hughes, Sébastien Vacher

**Affiliations:** CONIDIA, Parc d’activités en Chuel, Route de Chasselay, 69650 Quincieux, France; Institut de Génomique Fonctionnelle de Lyon (IGFL), CNRS UMR 5242, Ecole Normale Supérieure de Lyon, INRAE USC 1370, Université Claude Bernard Lyon 1, 69364 Lyon Cedex 07, France; CONIPHY, Parc d’activités en Chuel, Route de Chasselay, 69650 Quincieux, France; CJH SARL, 21 C Chemin de la Girotte, 33650 La Brede, France

**Keywords:** cytochrome *b*, downy mildew, complex III inhibitors, resistance, long and short reads sequencing, L201S, E203-DE-V204, E203-VE-V204

## Abstract

Downy mildew is caused by *Plasmopara viticola*, an obligate oomycete plant pathogen, a devasting disease for grapevine. To preserve plants from the disease, complex III inhibitors are among the widely used fungicides that specifically target the mitochondrial cytochrome *b* (cyt*b*) of the pathogen to block cellular respiration mechanisms. In French vineyard, *P. viticola* developed resistance against a first category of these fungicides, the Quinone outside inhibitors, by exhibiting a single amino acid substitution G143A in its cyt*b* mitochondrial sequence. Their usage was restricted and another kind of fungicides, Quinone inside inhibitors, targeting the same gene and highly effective against oomycetes, were used instead. Recently however, less sensitive *P. viticola* populations were detected after treatments with some inhibitors, in particular ametoctradin and cyazofamid. By isolating resistant single-sporangia strains of *P. viticola* to these fungicides, we characterized new variants in cyt*b* sequences associated with cyazofamid resistance: a point mutation (L201S) and more strikingly, two insertions (E203-DE-V204, E203-VE-V204). In parallel with classical tools, pyrosequencing and RT-PCR, we then benchmarked both short and long-reads NGS technologies (Ion Torrent, Illumina, Oxford Nanopore Technologies) to sequence the complete cyt*b* with the prospect to detect and assess the proportion of resistant variants of *P. viticola* at a natural population scale. Eighteen populations collected from French vineyard fields in 2020 were analysed: 12 show a variable proportion of G143A, 11 of E203-DE-V204 and 7 populations of the S34L variant that confers resistance to ametoctradin. Interestingly, long reads were able to identify variants, including SNPs, with confidence and detect a small proportion of *P. viticola* showing several variants along the same cyt*b* sequence. Altogether, NGS appear promising methods to evaluate pathogen resistance towards fungicides related to cyt*b* modifications at a population scale in the field. This approach could be rapidly a robust decision-support management tool for vineyard in future.

## Introduction

Fungal plant pathogen diseases can damage plant and crops, causing highly destructive impact in agriculture activities and food production. *Plasmopara viticola* is an obligate oomycete plant pathogen and the causal agent of downy mildew, the most devastating disease in grapevine. *P. viticola* infects all green organs of host plant and has an alternating life cycle between sexual overwintering phase and asexual multiplication during the growing season, causing primary and secondary infection cycles respectively [1–6]. Currently management strategies of downy mildew include multi-site fungicides like copper-based fungicides, such as Bordeaux mixture and the dithiocarbamates, as preventive treatment. Then, single-site fungicides, such as phenylamides (e.g. metalaxyl), QoI (Quinone outside Inhibitors; e.g. azoxystrobin), carboxylic acid amides (CAA; e.g. mandipropamid), and more recently, complex III inhibitors QiI’s (Quinone inside Inhibitors; e.g cyazofamid and amisulbrom), are often introduced in management program (see Fungicide Resistance Action Committee – FRAC - Code List 2021 for details [7]). Complex III inhibitors target the mitochondrial cytochrome *b* (cyt*b*) protein and block cellular respiration mechanisms [8].

The mitochondrial respiratory chain consists of multifunctional, oligomeric membrane enzyme complex. Cytochrome *bc1* complex (complex III) is a key enzyme in mitochondrial electron transport chain. Cyt*b* is a subunit of complex III that catalyses the transfer of electrons from ubiquinol to cytochrome *c* leading to protons translocation and energy transduction. The cyt*b* protein contains eight transmembrane helices encoded by the cyt*b* gene. The widely used complex III inhibitor’s fungicides target sites on these helices with different strategies [8,9].

The fungicides known as QoI’s block mitochondrial respiration by binding to Qo site [9,10]. QoI fungicide-resistant isolates were detected in field populations of many plant pathogens such as *Erysiphe necator* [11,12], *P. viticola* [13], *Alternaria sp*. [14,15], *Mycosphaerella graminicola* [16,17] and many others [7]. In French vineyard, downy mildew populations (*P. viticola)* have developed resistance to QoI’s fungicides by a single amino acid substitution G143A in cyt*b* [9,13,18,19]. The use of this single-target fungicide group was restricted after the widespread of the resistance.

The fungicides known as QiI’s inhibit the reduction of quinol in Qi site close to the mitochondrial matrix. These fungicides are highly effective against oomycetes and play an important role in downy mildew management programs nowadays. Recently however, less sensitive *P. viticola* populations to ametoctradin (QoSI fungicide, Quinone outside Stigmatellin binding sub-site Inhibitor; FRAC Code List 2021 [7]) and cyazofamid (QiI fungicide), were detected in vineyard [20,21]. The origin of this resistance was unknown with different possible hypotheses. The specific resistance could result of new modifications in the cyt*b P. viticola* sequence, such as S34L that is predicted to destabilise the fungicide binding in the case of ametoctradin [22], or potentially new mutations never described yet. Alternatively, or in combination, and although mutations in the inhibitor binding sites within cyt*b* represent most cases of reported resistance, it is known that other mechanisms may induce resistance to some fungicides ([22] for review), in particular by the activation of a mitochondrial alternative oxidase (AOX). This mechanism of alternative respiration pathway was already observed in collected *P. viticola* populations that were not submitted to QoSI (ametoctradin) and QiI’s (cyazofamid) pressure selection [20,21].

To better characterize the origin of the new resistance observed in the vineyard, the aim of this study was to isolate sensitive and resistant single-sporangia strains to ametoctradin and cyazofamid by biological tests and analyse the resistant strains by molecular approaches. Leaf disc sensitivity bioassay was performed on isolated strains to investigate cross-resistance between QoI, QiI and QoSI fungicides. Cyt*b* sequences of the isolated single-sporangia strains were sequenced by Sanger and analysed to characterize new possible molecular mechanisms of resistance to these fungicides in *P. viticola*. To go further and analyse cyt*b* polymorphism at a natural population scale, NGS technologies were first benchmarked to sequence the complete cyt*b* from single-sporangia strains and then applied to analyse field populations of *P. viticola*. Still relatively poorly present in studies associated to plant pathogen resistance to fungicide and targeting usually full pathogen genomes [23,24], both short-read (Ion Torrent and Illumina) and long-read (Oxford Nanopore Technologies, ONT) technologies were tested and compared in our study. They revealed to be efficient to identify and monitor cyt*b* variants in natural populations but also promising to become a robust tool for decision-support in management of fungicide resistance, potentially directly in the field, in near future.

## Materials and methods

### *P. viticola* populations and culture conditions

Downy mildew infected leaves were randomly collected from vineyards of different France regions in 2016, 2017, 2018 and 2020. Sampling was carried out in vineyards with fungicide protection programs including or not complex III inhibitors. More than 50 leaf discs surrounding infected lesion (oil spot) per sample were prepared from collected leaves. Leaf discs were placed onto Petri dish, washed with distilled water and dried at room temperature. After 24h incubation, new sporangia of downy mildew were collected in sterile water to inoculate decontaminated fungicide-free leaves from grape cultivar *Vitis vinifera* cv. Cabernet-Sauvignon. Inoculated leaves were incubated in Petri dishes for 7 days at 22°C with a 16 hours of light:8 hours of dark photoperiodic lighting. Freshly produced sporangia were harvested to inoculate testing leaf discs.

### Chemical fungicides

Commercial formulations of ametoctradin (SNOOKER, concentrated solution containing 200g active ingredient (AI) L^-1^, BASF, France) and cyazofamid (RANMAN TOP, concentrated suspension containing 160g AI L^-1^, ISK Biosciences Europe, France) were tested. The fungicide formulations were dissolved in sterile distilled water. Stock solutions were stored at 4°C in the dark.

### Resistant single-sporangia isolates and cross-resistance assay

Leaf discs with sporulating colonies at 1mg.L^-1^ of cyazofamid or ametoctradin with 100mg.L^-1^ of SHAM (Salicylhydroxamic acid, Sigma-Aldrich, France), as inhibitor of alternative respiration (AOX), were used to isolate resistant strains of *P. viticola*. Collected populations showing high level of non-specific resistance on fungicide without SHAM were also used to isolate AOX-strains. Thus, monitoring of sensitivity assays using fungicide, alone or with SHAM, was used to select strains developing resistance involving either AOX respiration (non specific resistance) or cyt*b* target mutation mechanism. Indeed, strains showing sporulation on leaf discs with both fungicide and SHAM had specific resistance mechanism, probably inducing cyt*b* target modification. In other hand, strains growing only on fungicide without SHAM involve AOX activity.

To investigate cross-resistance, *in vitro* sensitivity of these strains to ametoctradin and cyazofamid, applied alone or mixed with SHAM, was measured with increasing concentrations of each fungicide (0.01mg.L^-1^ to 100mg.L^-1^). Ten discs were analysed for each condition and assays for each isolate were repeated three times per fungicide concentration. After 12 days, individual leaf discs were evaluated for disease incidence and sporulation rate.

### Total DNA extraction and PCR amplification

Single-sporangia strains of *P. viticola* growing leaf discs 7 days post inoculation were used as starting material for DNA extraction and PCR amplification. Total DNA was extracted using the Nucleospin® plant II kit (Macherey-Nagel GmbH & Co., Düren, Germany) according to the manufacturer’s recommendations. The same approach was used for natural populations collected from field, except that DNA was extracted from freshly inoculated leaves as previously described.

PCR amplification of 1kb fragment of cyt*b* gene was carried out in 25μL reaction mixtures with 30ng of total genomic DNA, set of forward (5’-TGAACCTGTAAATTTAGCACAACAA-3’) and reverse (5’-ACAGGACATTGACCAACCCA-3’) primers (0.3μM) and 1X premixed Phusion Flash High-Fidelity PCR Master Mix (Thermo Scientific, France). Amplifications were carried out in a thermal cycler LifeEco (BIOER Technology, France) using the following PCR program: initial denaturation at 94°C for 1min followed by 35 cycles at 95°C for 30s, 60°C for 1min, and 72°C for 1 min and a final extension at 72°C for 10min. Amplification products were then subjected to direct Sanger sequencing, using the same primers as for PCR amplification (GATC Biotech, Germany). To analyse cyt*b* gene partial sequence and investigate point mutations in resistant strains, the new sequences were aligned with Jalview [25] against a *P. viticola* cyt*b* sequence used as reference (accession number DQ459459.1) and against sequences of sensitive strains.

### Pyrosequencing assay design for cyt*b* alleles quantification

Pyrosequencing technology was used to investigate allele’s frequencies of modifications detected in cyt*b* genes amplified from different samples: field isolated strains (selected single-sporangia strains) and samples from natural populations (mixed sporangia collected in the field). PCR reactions for pyrosequencing were performed in a final volume of 50µl containing 30ng genomic DNA, 12.5µl of PyroMark PCR Master Mix (Qiagen, France), 2.5µl of CoralLoad concentrate (Qiagen, France), 5µM of CONIPHY designed reverse and forward primers (*P. viticola* PyroID kit, CONIPHY, France) using PyroMark Assay Design version 2.0 (Qiagen, France). Target region of cyt*b* gene with the insertion of nucleotides was amplified using the following program: 95°C for 15min, followed by 45 cycles (94°C for 30s, 60°C for 30s and 72°C for 30s) and a final DNA extension at 72°C for 10min (following manufacturer recommendations). PCR products (3µl of the initial reaction mixture) were separated on a 1.2% agarose gel stained with SYBR^®^ Safe (Invitrogen™, France). Gel production and electrophoresis were conducted using TAE buffer (40mM Tris-acetate pH 8.0, 1mM EDTA). Pyrosequencing reactions were performed in a PyroMark Q48 Autoprep instrument (Qiagen, France) using PyroMark^®^ Q48 Advanced Reagents kit (Qiagen) with 3µl of PyroMark Q48 magnetic beads (Qiagen) and 10µl of biotinylated PCR products following manufacturer’s instructions. The sequencing primers provided in dedicated PyroID kit (CONIPHY, France) were used to detect E203-DE-V204 and L201S or E203-VE-V204. For validation of assays, 5 replicates of total DNA extracted from sensitive and resistant single sporangia isolates were analysed. Allele frequencies were estimated using PyroMark software.

### Cyt*b* PCR amplification and sequencing by short and long reads NGS technologies

NGS technologies were explored to characterize cyt*b* variants from sensitive or resistant single-sporangia isolated strains (7 DNA extracts tested). After this validation step, NGS were used to detect the presence and proportion of sensitive strains among natural populations collected in French vineyard in 2020 (18 DNA extracts tested). This was done by amplifying the complete cyt*b* from the total DNA extracted and by sequencing the amplicons by short and long-read technologies.

For all the 25 samples analysed, complete cyt*b* was amplified using two different strategies: in 5 overlapping fragments (302 to 375 bp long) to perform short-read sequencing (Illumina or Ion Torrent), and in a single fragment (1457 bp long) to perform long-read sequencing (ONT).

Five primer pairs were designed by Ion AmpliSeq Designer (Thermofisher Scientific) using DQ459459.1 as a reference. For short-read approaches, the five overlapping fragments were amplified separately in 25µl in a mix including 2X TaqMan™ Environmental Master Mix 2.0 (Applied Biosystems), 1µl of each primer at 10nM and 10ng of genomic DNA. PCR amplifications were carried out in a Veriti Thermocycler (Thermofisher Scientific) using the following PCR conditions: initial denaturation at 94°C for 5s followed by 35 cycles at 94°C for 30s, 55°C to 65°C (depending on primers set) for 45s, and 72°C for 1min and a final extension at 72°C for 7min. The five amplicons were then pooled for each sample in an equimolar manner. For long-read approach, long-range PCR was performed to amplify the complete gene with the same mix composition as above except that the LongAmp HotStart Taq 2X Master Mix (New England Biolabs) was used instead. The primers used correspond to the most extreme forward and most extreme reverse primers designed above. The PCR program was the following: initial denaturation at 94°C for 10min followed by 40 cycles at 94°C for 30s, 60°C for 45s, and 65°C for 2min and a final extension at 65°C for 7min.

For Ion Torrent, the barcoded libraries were built from pooled amplicons by a ligation protocol using the Ion Xpress™ Plus Fragment Library Kit and following the recommendations of the manufacturer. Pooled libraries were sequenced in SE 400bp on a 318 Ion chip using a PGM sequencer (Thermofisher Scientific). For Illumina and ONT, fusion primers (i.e. containing partial adapters sequences specific of respectively Illumina or ONT) were used for the first PCR. In both cases, a second PCR was performed to add indexes/barcodes and to complete the libraries construction. Illumina barcoded libraries were sequenced with a Reagent kit v2 in PE 2×250bp on a MiSeq sequencer. Nanopore libraries were finalized and barcoded with the PCR Barcoding kit protocol (SQK-PBK004, ONT) and sequenced on Flongle flow cells (FLO-FLG001, ONT) with Mk1C or MinION coupled with MinIT. High accuracy basecalling was performed for all Nanopore runs with Guppy version 3.2.9 or 4.3.4.

Short and long raw reads datasets were deposited to SRA database under BioProject accession N°XXXXXX.

### NGS data analyses

Ion Torrent reads were analysed using the AmpliSeq design above and DQ459459.1 as a reference. Torrent Variant Caller (TVC) plugin proposed by Thermofisher Scientific in the Torrent Suite Software was launched with a generic configuration for somatic and low stringency parameters. In this configuration, 2000 reads are considered to characterize new variants and low frequency detection is optimized (usually detection is not given below 5%). Illumina reads were analysed with Galaxy Europe (https://usegalaxy.eu) or with our own instance. Roughly, for each sample, R1 and R2 reads were paired and then sorted by type of fragment. Five thousand reads were sampled for each of the 5 fragments and mapped with minimap2 [26] on the DQ459459.1 reference. The Bayesian genetic variant detector FreeBayes [27] was used (option simple diploid calling with filtering and coverage) to find polymorphisms and VCFlib [28] used to extract the variable positions in a table format with reads count.

Nanopore reads were analysed with a dedicated pipeline designed in command lines. Only reads higher than Q7 were considered. Five thousand reads were sampled for each sample and mapped on the DQ459459.1 reference with minimap2. Six regions or positions identified by this study as variable were searched with either a BLAST [29] approach or a Nanopolish (https://github.com/jts/nanopolish) analysis. Frequency at each position was estimated by counting the number of reads with the variant versus the total reads number.

To identify complete cyt*b* sequence with multiple polymorphisms, each single polymorphism was searched independently among the previous 5000 long reads sampled. All unique reads containing at least one polymorphism were then considered as a whole and proportion of reads containing one or multiple variants along the same read were computed. R package “ggvenn” was used to draw Venn diagrams and to obtain percentage of reads corresponding to each possible combination. Presence of several polymorphisms along the same read was checked by aligning read against the reference and confirmed by eye on randomly picked reads.

## Results and discussion

### Sensitivity of field isolates to ametoctradin and cyazofamid

Many downy mildew field populations with low sensitivity to ametoctradin and cyazofamid at different levels were observed in French vineyards since 2016. Resistant and sensitive populations were selected to generate single-sporangia isolates. More than 70 single-sporangia strains were isolated including 41 resistant to cyazofamid (QiI), 5 to ametoctradin (QoSI), 7 to pyraclostrobin (QoI, data not shown), 15 to ametoctradin and pyraclostrobin, and 3 strains showing non-specific resistance with AOX expression. Further investigation on cross resistance and genotypic characterization was conducted on a panel of 22 strains listed in Table 1.

**Table 1:** MIC (Minimum Inhibitory Concentration) values for ametoctradin and cyazofamid of *P. viticola* field isolated strains and detected mutations or insertions in cyt*b* gene compared to the reference DQ459459.1. A dash indicates no modification, a cross indicates detection of the variant. R: resistant; S: Sensitive, indicate resistance state when AOX mechanism is blocked by SHAM and probably results from cyt*b* modifications in most part.

| *P. viticola*<br>strains | Ametoctradin (QoSI) |  |  | Cyazofamid (QiI) |  |  | Modification of *cytb* sequence observed |  |  |  |  |
| --- | --- | --- | --- | --- | --- | --- | --- | --- | --- | --- | --- |
|  | Alone | +SHAM |  | Alone | +SHAM |  | G143A | S34L | E203-DE-V204 | E203-VE-V204 | L201S |
| CONI-01 | 0.3 | <0.1 | S | <0.1 | <0.1 | S | - | - | - | - | - |
| CONI-04 | >100 | 0.3 | S | 30 | <0.1 | S | - | - | - | - | - |
| CONI-02 | 0.3 | 0.3 | S | <0.1 | <0.1 | S | X | - | - | - | - |
| CONI-03 | 0.3 | 0.3 | S | <0.1 | <0.1 | S | X | - | - | - | - |
| CONI-11 | >100 | 0.3 | S | 30 | <0.1 | S | X | - | - | - | - |
| CONI-12 | >100 | 0.3 | S | 30 | <0.1 | S | X | - | - | - | - |
| CONI-20 | >100 | 100 | **R** | <0.1 | <0.1 | S | - | X | - | - | - |
| CONI-22 | >100 | 100 | **R** | <0.1 | <0.1 | S | - | X | - | - | - |
| CONI-06 | >100 | 100 | **R** | <0.1 | <0.1 | S | X | X | - | - | - |
| CONI-13 | 100 | 100 | **R** | <0.1 | <0.1 | S | X | X | - | - | - |
| CONI-05 | 1 | 1 | S | 100 | 30 | **R** | - | - | X | - | - |
| CONI-07 | 1 | 0.3 | S | 100 | 30 | **R** | - | - | X | - | - |
| CONI-15 | 1 | 0.3 | S | 100 | 30 | **R** | - | - | X | - | - |
| CONI-16 | 1 | 0.3 | S | 100 | 30 | **R** | - | - | X | - | - |
| CONI-08 | >100 | 0.3 | S | 100 | 30 | **R** | - | - | X | - | - |
| CONI-09 | >100 | 0.3 | S | 100 | 30 | **R** | - | - | X | - | - |
| CONI-10 | >100 | 0.3 | S | 100 | 30 | **R** | - | - | X | - | - |
| CONI-17 | 1 | 0.3 | S | 100 | 30 | **R** | - | - | X | - | - |
| CONI-31 | >100 | 0.3 | S | 30 | 30 | **R** | - | - | - | - | X |
| CONI-38 | 10 | 0.3 | S | 30 | 30 | **R** | - | - | - | - | X |
| CONI-39 | >100 | 0.3 | S | 30 | 10 | **R** | - | - | - | X | - |
| CONI-41 | 0.3 | 0.3 | S | 30 | 10 | **R** | - | - | - | X | - |

### Molecular analysis of field isolates *P. viticola* cyt*b* gene Confirmation of S34L substitution in cytochrome *b* conferring resistance to ametoctradin

According to their sensitivity to ametoctradin, growth of 4 strains of *P. viticola* (CONI-06, CONI-13, CONI-20, CONI-22) was inhibited at 100 mg.L^-1^ of ametoctradin applied alone or mixed with SHAM. These strains exhibited high level of resistance (Resistance Factor = 1000) to ametoctradin compared with sensitive strain CONI-01 (MIC <0.1 mg.L^-1^). These strains are sensitive to cyazofamid (<0.1 mg.L^-1^) applied alone or with SHAM (Table 1). Loss of sensitivity in these strains seems to be caused by target modification affecting specifically ametoctradin mode of action. Analysis of cyt*b* gene sequences of these strains, obtained by Sanger sequencing, reveals the presence of single nucleotide mutation of cytosine to thymine at position 101 (TCA → TTA). This mutation in cyt*b* gene leads to substitution of amino acid serine with leucine at position 34 (S34L) in ametoctradin resistant *P. viticola* isolates (Fig 1A). In other hand, no cross resistance was observed in strains carrying S34L substitution with cyazofamid. Among strains with G143A substitution, conferring resistance to QoI fungicide, cyt*b* gene sequence of two isolates (CONI-06 and CONI-13) contains both S34L and G143A substitutions (Table 1). Tested with leaf discs bioassays, these strains are resistant to both pyraclostrobin (QoI) and ametoctradin (data not shown). Ametoctradin has been first described as QoSI inhibitor acting on the Qo site [30]. However, serine S34 of *P. viticola* cyt*b* is located in the Quinone inside site (Qi) suggesting that ametoctradin inhibits mitochondrial respiration by interacting with complex III in Qi site. The results suggest that the mode of action of ametoctradin in Qi site is different from that of cyazofamid. They confirm recent investigation on the binding mode of ametoctradin with its target site in cytochrome *bc1* complex showing that the anti-oomycete fungicide ametoctradin is able to interact with both Qo and Qi sites [20,22,31].

**Fig 1.**
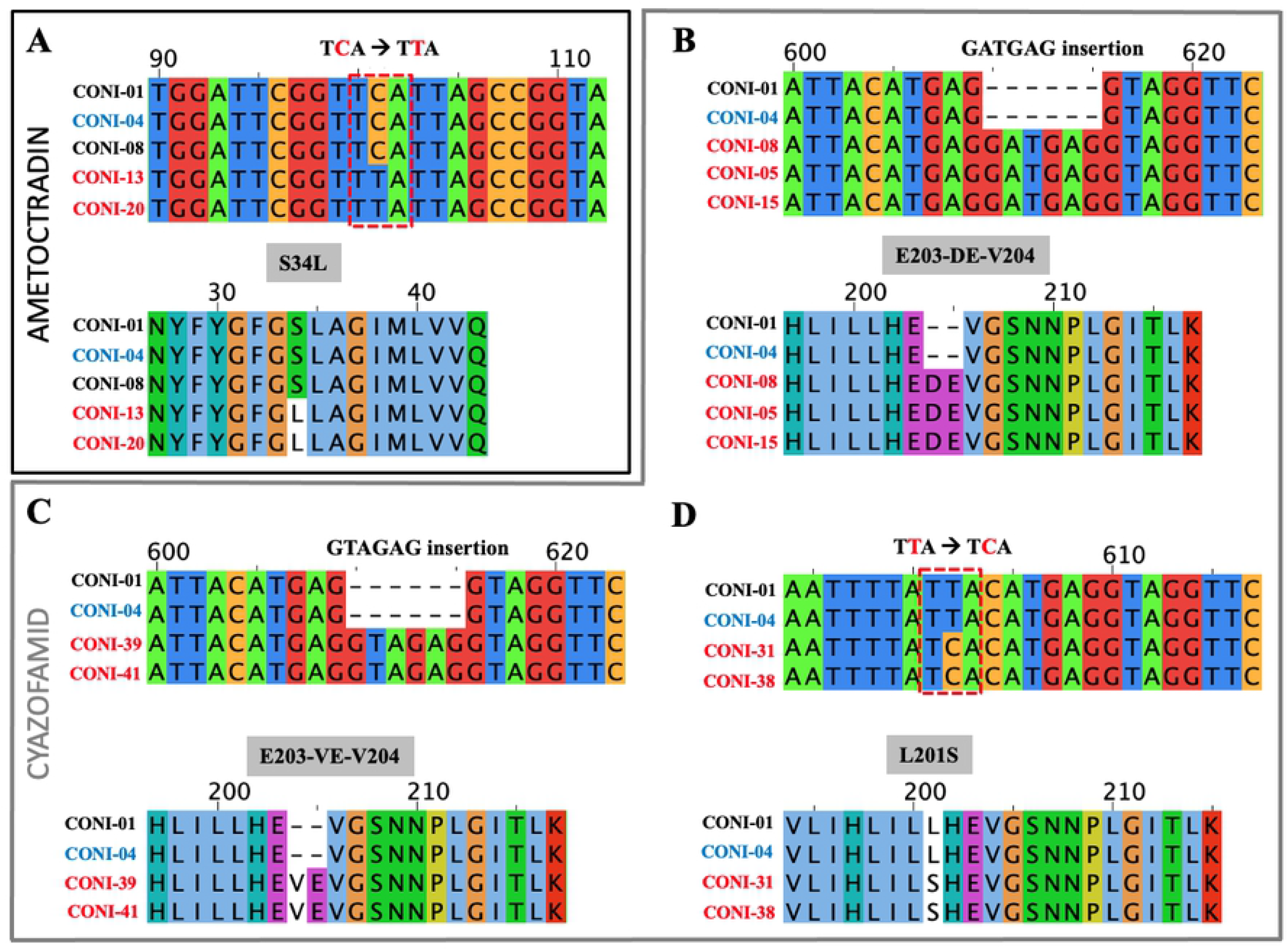
Alignment of cyt*b* partial sequences of cyazofamid and ametoctradin *P. viticola* resistant isolates. In black: sequence of sensitive strain. In red: sequences of strains resistant to cyazofamid or ametoctradin accordingly. In blue: AOX strain. Sequences of strains with S34L substitution **(A)**, GATGAG insertion **(B)**, GTAGAG insertion **(C)** and L201S substitution **(D)**.

### L201S substitution and two different 6bp insertions (E203-DE-V204 and E203-VE-V204) in cyt*b* coding gene confer resistance to cyazofamid

Sensitivity bioassays revealed 41 isolates of *P. viticola* resistant to cyazofamid applied alone or mixed with SHAM at a discriminating dose (1mg.L^-1^). Further analysis of 12 strains reveals that *P. viticola* development of 10 isolates was inhibited at 30mg.L^-1^ and 2 isolates (CONI-39 and CONI-41) at 10mg.L^-1^ of cyazofamid+SHAM compared to sensitive strain CONI-01 (<0,1 mg.L^-1^) (Table 1). The cyt*b* sequences of 31 of these isolated resistant strains show a 6 nucleotides insertion, GATGAG, compared to cyazofamid-sensitive strains and the reference DQ459459.1. This short sequence insertion leads to protein modification with two additional amino acids (E203-DE-V204) (Fig 1B). A second 6 nucleotides insertion, GTAGAG, is observed in two other strains leading to another amino acid modification E203-VE-V204 (Fig 1C). Finally, cyt*b* sequences of 2 isolated strains, referenced CONI-31 and CONI-38, inhibited at 30mg.L^-1^ of cyazofamid+SHAM, contain a single nucleotide mutation at position 602 (TTA→TCA) inducing amino acid substitution L201S in cyt*b* protein (Fig 1D). These three variants, all occurring in the same short region of the cyt*b* gene, confer resistance to cyazofamid but no cross resistance with ametoctradin. To our knowledge, fungicide resistance caused by target modification occurred only with single nucleotide modification (SNP) in almost described plant pathogens and fungicides single-site mode of action [7,32]. Insertion of short sequence in gene coding fungicide-target, and probably protein conformational modification, without altering protein function, could be a new way to bypass the fungicide action. Interestingly, similar amino acid modification was introduced in yeast cyt*b* mutant model and no effect was observed on growth of yeast. Comparative protein structure modelling suggests that the two amino acid insertion E203-DE-V204 interfere with cyazofamid binding to cyt*b* in *P. viticola* [33]. Description of six sequence variants in *P. viticola* cyt*b* gene that are associated with resistance to cyazofamid, ametoctradin or QoI fungicides, identified in this study or coming from the literature, are listed in Table 2 and were targeted in the further analyses.

**Table 2:** Description of *P. viticola* variants of cyt*b* gene and associated resistance to cyazofamid, ametoctradin or QoI fungicides considered in this study. DQ459459.1 is used as the reference.

| Variants | AA modifications | Reference | Variant | Type | Resistance | References |
| --- | --- | --- | --- | --- | --- | --- |
| Variant 1 | E203-DE-V204 | - | GATGAG | Insertion (6bp) | Cyazofamid<br>QiI fungicide | [20], this study |
| Variant 2 | E203-VE-V204 | - | GTAGAG | Insertion (6bp) | Cyazofamid<br>QiI fungicide | This study |
| Variant 3 | L201S | T | C | Substitution | Cyazofamid<br>QiI fungicide | This study |
| Variant 4 | S34L | C | T | Substitution | Ametoctradin<br>QiSI fungicide | [20, 21], this study |
| Variant 5 | F129L | T | A/G/C | Substitution | QoI fungicides | FRAC 2021 |
| Variant 6 | G143A | G | C | Substitution | QoI fungicides | FRAC 2021, this study |

### Pyrosequencing quantification of L201S substitution and the two insertions in *P. viticola* cyt*b* gene

Two pyrosequencing-based methods were developed to quantify the frequency of the 3 variants in samples. The frequencies of E203-DE-V204 and L201S were evaluated simultaneously in a first assay (Supplementary Fig 1) and the frequency of E203-VE-V204 in a second assay (Supplementary Fig 2). Pyrosequencing assays were validated on DNA extracted from single sporangia cyazofamid-resistant isolates. Pyrosequencing assays on DNA extracted from single sporangia showed that cyazofamid-resistant isolates had between 96 to 98% insertions E203-DE-V204, E203-VE-V204 or substitution L201S, corroborating Sanger analysis (Fig 1) and sensitivity assays (Table 1). Finally, to evaluate the background noise, the two methods were applied to the sensitive wild-type strain (CONI-01) that present no sequence variants. In both cases, low frequencies (2-8%) of variants were observed (Supplementary Fig 1 and Fig 2) allowing to fix a threshold for accurate detection. For further analyses, only populations with variant frequency higher than 10% were considered as significantly detected by pyrosequencing. Applied to DNA extracted from natural populations collected in the field, only variant E203-DE-V204 was detected at different proportion using pyrosequencing analyses. Frequencies of variant E203-DE-V204 below 5%, or close to background noise, were measured in five populations sensitive to cyazofamid (100% efficacy; Table 3). Eleven *P. viticola* field populations contain variant E203-DE-V204 with frequencies ranging from 10 to 96%. Both observations, on single sporangia isolates and field natural populations, show significant correlation between genetic profiling by pyrosequencing and phenotypic characterization with *in vitro* biological assays. Variants E203-VE-V204 and L201S are not detected in the natural populations tested (Table 3), suggesting that the insertion E203-DE-V204 could be predominant in the field and probably at the origin of downy mildew populations resistant to cyazofamid in French vineyards.

**Table 3:** Characterization of fungicide resistance of 18 natural populations of *P. viticola* collected in French vineyards in 2020. Biological tests and associated proportion of *P. viticola* sequence variants observed from cyt*b* gene estimated by RT-PCR, pyrosequencing and NGS sequencing are given. For the biological tests, value of 100% indicates no resistance measured in the population for the fungicide, and 0% means totally resistant. Sequence cyt*b* variants considered are described in Table 2. Only variants E203-DE-V204, S34L and G143A were detected in the field populations analysed and are given. All values are expressed in %. A dash indicates non detected or value below limit detection (threshold of 5% considered for NGS).

| Samples |  | Biological test on leaf discs (% efficacy) |  | Cytb allele frequency quantification |  |  |  |  |  |  |  |  |
| --- | --- | --- | --- | --- | --- | --- | --- | --- | --- | --- | --- | --- |
| Field population | Location French department | Ametoctradin (1mg/l) + SHAM (100mg/l) | Cyazofamid (1mg/l) + SHAM (100mg/l) | RT-PCR |  | Pyrosequencing<br>% E203-DE-V204 | NGS sequencing |  |  |  |  |  |
|  |  |  |  | % G143A | % S34L |  | FreeBayes / Illumina (Short reads) |  |  | BLAST / ONT (Long reads) |  |  |
|  |  |  |  |  |  |  | % G143A | % S34L | % E203-DE-V204 | % G143A | % S34L | % E203-DE-V204 |
| CONI-P1 | 33 | 100 | 35 | - | - | 52 ± 0.7 | - | - | 51 | - | - | 48 |
| CONI-P2 | 30 | 100 | 100 | - | - | 3 ± 0.4 | - | - | - | - | - | - |
| CONI-P3 | 47 | 100 | 100 | - | - | 4 ± 1.3 | - | - | - | - | - | - |
| CONI-P4 | 13 | 100 | 100 | - | - | 4 ± 1.6 | - | - | - | - | - | - |
| CONI-P5 | 51 | 98 | 0 | 8 | - | 81 ± 1.1 | 8 | - | 81 | 8 | 7 | 81 |
| CONI-P6 | 51 | Not tested | 97 | 94 | - | 9 ± 0.4 | 93 | - | - | 94 | - | 6 |
| CONI-P7 | 51 | Not tested | 82 | 63 | - | 33 ± 0.9 | 60 | - | 30 | 64 | - | 28 |
| CONI-P8 | 51 | Not tested | 60 | 46 | - | 31 ± 0.7 | 43 | - | 30 | 46 | - | 28 |
| CONI-P9 | 51 | Not tested | 23 | - | - | 96 ± 1.1 | - | - | 95 | - | - | 95 |
| CONI-P10 | 17 | 100 | 69 | 59 | - | 37 ± 1.1 | 55 | - | 35 | 57 | - | 36 |
| CONI-P11 | 33 | 100 | 85 | - | - | 25 ± 1.1 | 13 | - | 23 | 13 | - | 22 |
| CONI-P12 | 33 | 100 | 100 | 56 | - | 6 ± 0.9 | 47 | - | - | 48 | - | - |
| CONI-P13 | 17 | 47 | 67 | 73 | 88 | 10 ± 0.9 | 69 | 94 | 5 | 70 | 93 | 5 |
| CONI-P14 | 17 | 40 | 100 | 74 | 79 | 4 ± 1.3 | 82 | 91 | - | 85 | 92 | - |
| CONI-P15 | 32 | 65 | 85 | - | - | 7 ± 0.9 | 6 | 9 | - | 6 | 12 | - |
| CONI-P16 | 33 | 55 | 57 | - | 18 | 85 ± 2.4 | - | 10 | 85 | - | 14 | 84 |
| CONI-P17 | 33 | 52 | 88 | 12 | 49 | 15 ± 1.1 | 8 | 57 | 12 | 9 | 59 | 12 |
| CONI-P18 | 51 | 57 | 80 | 39 | 63 | 13 ± 2.9 | 43 | 64 | - | 43 | 64 | - |

**Fig 2.**
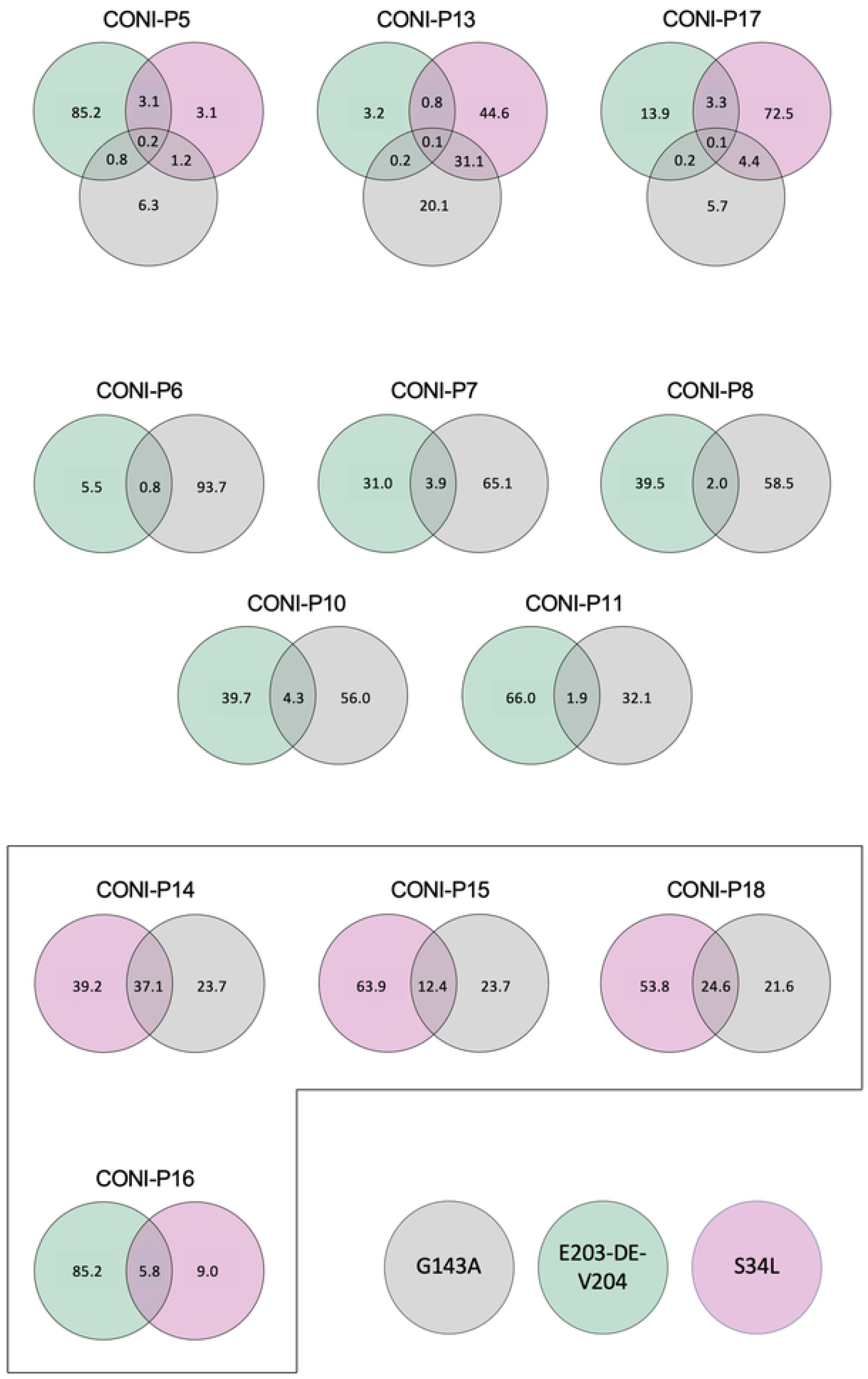
Cyt*b* sequences coming from field *P*.*viticola* populations can exhibit several modifications correlated to resistance to cyazofamid and ametoctradin. For each population where at least two different variants were detected (Table 3), and considering only ONT reads with variants, the percentage of each type of variant (E203-DE-V204, S34L or G143A; Table 2) observed in the same read is indicated in the Venn diagram. Framed diagrams indicate populations with more than 5% reads exhibiting two different variants along the same sequence.

Molecular characterization tools, including real-time PCR (RT-PCR), are widely used in detection and quantification of fungicide resistance in plant pathogens [11,21,34–38]. In this particular case however, RT-PCR assay design failed to analyse the two insertions of six nucleotides observed in *P. viticola* cyt*b* gene, their detection relying on the pyrosequencing approach only. Allele quantification using pyrosequencing technology has been used for SNP detection and quantification of QoI and CAA fungicides resistance in *P. viticola* [39,40]. In other hand, target site modification related to succinate dehydrogenase inhibitors (SDHI) resistance in *Pyrenophora teres* were identified using pyrosequencing [41]. Fields monitoring of G143A substitution in *Cercospora sojina* [42] or resistance to DMI and SDHI of *Ramularia collo-cygni* populations [43] have been assessed by pyrosequencing. Pyrosequencing method is a powerful alternative method allowing detection and quantification of single nucleotide polymorphism (SNP) but also, as reported in this study, that permits the detection of nucleotides insertion or deletion to assess gene modification in fungal fungicide resistance. Pyrosequencing and RT-PCR methods are accurate and less time-consuming approaches than biological assays. However, these methods are only relevant in the case of fungicide resistance mechanisms when modifications of the target are known [38]. Moreover, many reactions can be needed for a single sample if various mutations associated to fungicide resistance have to be assessed, such as G143A, S34L and L201S for example.

Considering the limits of the two previous methods, we explored others approaches to identify new variants or to assess the presence of known cyt*b* variants and their proportion at a population scale. In this context, more recent development of new sequencing technologies, such as short and long reads NGS, are promising and open opportunities to improve quality and sensitivity of molecular detection of pathogen resistance to fungicide. For example, Whole Genome Sequencing (WGS) is already largely used to track antimicrobial resistance [44], sometimes combining short and long reads approaches [45]. In medical applications, ONT long reads generated in real-time raise hope for very rapid diagnostic to identify pathogens and antibiotic resistance [46]. ONT reveals an interesting tool for plants surveillance to detect plant virus routinely [47]. Finally, large-scale genomics studies of plant disease resistance were possible thanks to these high-throughput and cost-effective tools to clarify the interactions between legumes and pathogens [48]. At a gene scale, only few studies already reported successful cyt*b* sequencing by short-read NGS method for the purpose of pathogen resistance to fungicides studies. It was the case for example to detect and characterize resistance of wheat pathogen (*Zymoseptoria tritici*) to QoI [49]. However, the NGS technologies have been poorly explored in the literature for that specific purpose. We so decided to test short and long reads sequencing with the aim to develop new methods to obtain a more efficient and rapid characterization of new or known cyt*b* variants of *P. viticola* in a context of fungicide resistance and that would be adapted to large population scale monitoring.

### NGS analyses of cyt*b* genes from field *P. viticola* isolates Benchmarking of the NGS technologies to accurately detect variants from single-sporangia strains

The complete cyt*b* gene amplified from 7 single-sporangia strains DNA extracts was analysed by long and short reads sequencing to evaluate the reliability of detection for the 3 cyt*b* variants that we described with Sanger (Fig 1, Table 2). The wild type strain CONI-01 is sensitive and served as a control, the other strains were characterized as resistant and exhibit either variant E203-DE-V204, E203-VE-V204 or L201S. Both short-read (Ion Torrent) and long-read (ONT) approaches succeeded to detect the expected variant at high frequency (all values > 92% and up to 100%) in the expected samples (Supplementary Table 1). TVC tool used for Ion Torrent reads, that reports only variants with frequency higher than 5%, find successfully the specific variant with high values for quality detection. Because the variant E203-VE-V204 is a duplicate of 6bp of the original sequence that is inserted, two alternative mappings are possible (Fig 1). Few mapping artefacts were thus observed that could result in the slightly lower frequency detected by VTC for this variant (92%). For ONT long-reads, both pipelines (BLAST or Nanopolish) gave close, similar and accurate estimates of the expected variant (>95%). Low frequencies (below 2% for BLAST and <5% for Nanopolish) are however detected in nearly all samples for which the variants were not expected. This suggests that below this threshold, detection of a variant cannot be trusted. According to this preliminary analysis, we arbitrarily fixed to 5% the threshold of variant detection for any NGS methods (short or long reads) and can be considered as a limit of the sensitivity of the method. Of note, the observation of a low percentage of the non-expected variant in the single-sporangia strains could be explained in different ways: by the experimental error-rate (errors induced during amplification by PCR or linked to the sequencing step), by the sensitivity of the tools used to analyse reads, but could also be an indication of heteroplasmy. Heteroplasmy is a widespread, but relatively rare, phenomenon observed in many species consisting in the presence of different types of mitochondrial DNA in the same cell [21]. Heteroplasmy has been associated as a possible way for some species to rapidly adapt to fungicide and develop resistance [50,51]. In particular, heteroplasmy has been reported for the G143A variant associated with QoI resistance [19,52], including for *P. viticola* [21].

### Exploiting NGS technologies to detect proportion of variants in *P. viticola* field populations

The DNA extracted from 18 natural field populations of *P. viticola* collected in 2020 from French vineyards were used to amplify the complete cyt*b* in short fragments for Illumina sequencing or in a single long fragment for ONT sequencing. Both sequencing revealed striking similar results in the detection and proportion of each variant among samples with high correlation values and R^2^ > to 99% (Table 3). Two small differences were observed with the detection by ONT approach of two variants in two supplementary samples (variant S34L for CONI-P5 and variant E203-DE-V204 for CONI-P6). However, both appear in a very low proportion (7% and 6% respectively, Table 3), close to the 5% chosen as a threshold to detect variants.

Only 3 out of the 18 samples revealed to be sensitive to the fungicides tested (ametoctradin and cyazofamid), and for which no one of the six cyt*b* variants (Table 2) we were looking for were detected by NGS sequencing. The variants L201S, E203-VE-V204 and F129L were not observed in any samples suggesting that they could be rare. However, the 3 other variants, E203-DE-V204, S34L and G143A were detected. The variant G143, well known in the literature and first detected in French vineyard in 2003 [53] with several occurrence [13], is the most present, with a detection in 12 samples. Then, the variant E203-DE-V204 (Table 2) is observed in more than 50% of the samples analysed in this study, with 11 samples showing the modified cyt*b* in different proportion (5 to 95%). This insertion, that we first detected in 2016 [20], is thus observed in several populations collected 4 years later and in elevated frequencies (> to 25% in 7 samples; Table 3). However, these results are to be taken with caution given the reduced number of samples analysed and have to be confirmed on a larger scale study. Finally, the variant S34L is detected in 6/7 samples, in various proportion (7 to 94%). The same variant was previously reported in 2017 where 7 out of the 33 vineyards tested using allele-specific PCR assays were identified for S34L [21].

Results observed for NGS sequencing (short or long reads) are mostly in agreement with other molecular methods used independently to assess the presence of E203-DE-V204 variant by pyrosequencing, or the presence of G143A and S34L variants by RT-PCR. However, NGS reveal to be potentially more sensitive than RT-PCR by detecting variants at low frequency in CONI-P11 and CONI-P15. Results are also in agreement with biological tests performed to evaluate the resistance to ametoctradin (associated to variant S34L) or to cyazofamid (associated to variant E203-DE-V204).

### Long reads can detect combination of variants along the same cyt*b* sequence

Short reads sequencing (Illumina or Ion Torrent) is interesting because of the high quality of the reads (reads with QV>25) and the low error rate, error rate that is even lower in Illumina reads than in Ion Torrent ones [54]. Both technologies are thus ideal to identify new substitutions or modifications with confidence. However, if overlapping short reads can be used and assembled to reconstruct a longer sequence when no diversity is present in the sample, they cannot be used to accurately perform phasing or to associate modifications occurring on different fragments in different proportion. At contrary long reads from ONT have lower quality (reads with QV<12), making sometimes more difficult to identify substitutions at low frequency. However, the diversity among a population of long fragments can be assessed with confidence, allowing to clearly identify different haplotypes and their proportion on long sequences.

Taking advantage of these NGS specificities, we searched in our datasets if in some cases, some variants could be present in combination along the same cyt*b* sequence or if each variant correspond to different cyt*b* sequences. The variants detected being not amplified on the same short fragments, long reads only, i.e. complete cyt*b* sequences, were explored. Reads obtained for 12 populations exhibiting at least 2 types of variants were analysed in more details (Table 3, Fig 2). The same threshold of 5% was applied to consider as significant a proportion of reads exhibiting a combination of variants, leading to restrict the dataset to only 4 populations.

In the CONI-P16 sample, around 40% of the reads identified with the variant S34L, contain also the variant E203-DE-V204 (Fig 2). Very few reads (5.8%) were concerned by this combined pattern suggesting that only few strains of *P. viticola* could be double-resistant to cyazofamid and ametoctradin for now, or that the simultaneous presence of both cyt*b* modifications could be deleterious for *P. viticola*. In consequence, the probability to isolate putatively viable single-sporangia strains by the experimental process described above was extremely low, and thus the possibility to describe them by Sanger sequencing alone was very limited. On contrary, the ONT approach used in this study illustrates the ability of the method to detect very early the apparition of multiple variants strains in a field population by long read sequencing, with a deep sequencing coverage of the complete cyt*b* obtained after PCR amplification.

By searching possible association of variant E203-DE-V204 and variant S34L with variant G143A in the same cyt*b* sequence in all populations detecting at least two of them (Fig 2), one can observed than variant S34L and G143A are more often associated than any other combination. Both variants are SNPs regularly described in the literature since years as key point cyt*b* mutations associated to resistance in plant pathogens [7]. For the 3 *P. viticola* populations collected that only shared S34L and G143A (CONI-P14, CONI-P15 and CONI-P18), 12.4% to 37.1% of reads with a variant are reads where the two variants are present. This is also the case for CONI-P13 (31.2%), in which 3 different variants were observed. In 3 out these 4 populations, both variants reach high percentages (Table 3), and are detected at low frequencies only in CONI-P15. It should be noticed that the 4 populations are localized in different departments, from different French regions, suggesting that this event could have occurred independently several times. The combination of G143A and S34L observed on long-reads suggests the existence of possible viable strains resistant to multiple fungicides (QoI and Ametoctradin), and could be not so rare. This is confirmed by the 2 resistant single-sporangia strains we were able to isolate in this study that combine the two variants (CONI-06 and CONI-13; Table 1) and by a similar strain identified in a previous publication [21].

## Conclusion

This study provides an overview on the structure of collected downy mildew populations in French vineyard. In addition of well-known resistance of *P. viticola* to QoI fungicides, resistant strains to other complex III inhibitors were detected. *P. viticola* develop resistance to ametoctradin by target modification with substitution S34L. New type of target modification by short nucleotides insertion was first time detected and reported by us in cyazofamid-resistant population in 2016. This mechanism consists in two different insertions of six nucleotides in cyt*b* gene in Qi site. In other strains, amino acid substitution L201S also confers resistance to cyazofamid. No cross resistance was observed between cyazofamid and ametoctradin in isolated strains.

After identifying cyt*b* variants of *P. viticola* associated with resistance, we combined classical (biological test assays, pyrosequencing and RT-PCR) and new sequencing approaches to monitor the variants at a population scale. This is a first study that combined all these methods at the same time at this scale. Analysis of 18 natural populations collected from field in 2020 show that many resistant phenotypes to complex III inhibitors, exhibiting different genotypes, can co-exist in the same population. In particular, short reads revealed to be highly efficient to detect and characterize new variants, and all NGS technologies to detect various proportion of resistant strains in natural populations. The conclusions reached, strikingly in agreement between the different NGS approaches, are congruent in nearly most cases with the results generated by other molecular experiments using independent methods traditionally used for that purpose. Taken together, NGS approaches appear promising to monitor fungicide resistance linked to cyt*b* modifications or detect new putative resistant strains in natural populations of *P. viticola*, with a better sensitivity and a cost that will be reduced when mutualising many samples at the same time. This should help to detect earlier the emergence and the propagation of some resistance in vineyard against some fungicides more rapidly, in a larger geographic scale and with a larger number of samples, avoiding the use of less efficient chemicals.

Although mechanisms of resistance towards complex III inhibitors are complex and multiple, when they are associated to cyt*b* sequence modification, long-read ONT sequencing should especially be seen as an interesting tool to assess resistance of pathogens potentially directly in the field thanks to portability and small size of ONT sequencers, as it was done to identify plant viruses [55–57]. Because the sequence error rate is expected to drastically drop for long reads in a near future, it should be possible to better detect and more rapidly characterize new cyt*b* resistant variants in natural populations, even when present at low frequencies. If ONT could revolutionize in-field downy mildew diagnostics and biosurveillance in the future [58], ONT long reads could also be a robust decision-support tool in fungicide treatments in vineyard, and more generally, for all plant of agricultural interest where resistance is associated with sequence modification.

## Acknowledgments

We thank the Freiburg Galaxy Team for the use of the Galaxy platform Europe. We also thank Emmanuel Quemener for the maintenance of the Galaxy platform at the ENS of Lyon.

## Author contributions

Conceptualization: SC, CH, BG, SH and SV

Formal Analysis: SC, HS, BG, JD, SH

Funding Acquisition: SC, CH, BG, SH, SV

Investigation: SC, LB, PR, BG, CH, HS, LP

Project Administration: SC, CH, BG, SH, SV

Supervision: SC, CH, HS, BG, SH, SV

Writing – Original Draft Preparation: SC, BG, SH, SV

Writing – Review & Editing: SC, LB, JD, PR, BG, SH, CH, HS, LP, SV

## Supporting information

**Fig S1. Pyrograms showing allele quantification of E203-DE-V204 and L201S variants from sensitive and cyazofamid-resistant *P. viticola* strains**. Pyrograms show DNA extract of single sporangia isolates CONI-01 (A), CONI-16 (B) and CONI-38 (C). Nucleotides position from 11 to 24 represents the variable region to be analysed to detect insertion E203-DE-V204 (B). Nucleotide in position 28 concern quantification of L201S substitution (C).

**Fig S2. Pyrograms showing allele quantification of E203-VE-V204 variant from sensitive and cyazofamid-resistant *P. viticola* strains**. Pyrograms show DNA extract of single sporangia isolates CONI-01 (A) and CONI-39 (B). Nucleotides positions 8 to 10 represents the variable region to be analysed to detect insertion E203-VE-V204 (B).

**Table S1: Detection and frequency of E203-DE-V204, E203-VE-V204 and L201S variants observed with short-read (Ion Torrent) and long-read (ONT) sequencing of cytb gene amplified from single sporangia strains**. For Ion Torrent data, variants were searched without a priori with Torrent Variant Caller (somatic and low stringency parameters) and only variants above 5% are reported. For ONT data, two dedicated pipelines using either BLAST or Nanopolish were tested. The 3 variants were specifically targeted in all samples.

## References

1. Fröbel S, Zyprian E. Colonization of Different Grapevine Tissues by Plasmopara viticola—A Histological Study. Frontiers in Plant Science. 2019;10. Available: https://www.frontiersin.org/article/10.3389/fpls.2019.00951

2. Armijo G, Schlechter R, Agurto M, Muñoz D, Nuñez C, Arce-Johnson P. Grapevine Pathogenic Microorganisms: Understanding Infection Strategies and Host Response Scenarios. Front Plant Sci. 2016;7: 382. doi:10.3389/fpls.2016.00382

3. Kamoun S, Furzer O, Jones JDG, Judelson HS, Ali GS, Dalio RJD, et al. The Top 10 oomycete pathogens in molecular plant pathology. Molecular Plant Pathology. 2015;16: 413– 434. doi:10.1111/mpp.12190

4. Rossi V, Caffi T. The Role of Rain in Dispersal of the Primary Inoculum of Plasmopara viticola. Phytopathology®. 2012;102: 158–165. doi:10.1094/PHYTO-08-11-0223

5. Gessler C, Pertot I, Perazzolli M. Plasmopara viticola: A review of knowledge on downy mildew of grapevine and effective disease management. Phytopathologia Mediterranea. 2011;50: 3–44. doi:10.14601/Phytopathol_Mediterr-9360

6. Rossi V, Giosuè S, Caffi T. Modelling the dynamics of infections caused by sexual and asexual spores during Plasmopara Viticola epidemics. Journal of Plant Pathology. 2009;91: 615–627.

7. FRAC | Home. [cited 28 Mar 2022]. Available: https://www.frac.info/

8. Huang L, Cobessi D, Tung EY, Berry EA. Binding of the Respiratory Chain Inhibitor Antimycin to the Mitochondrial bc1 Complex: A New Crystal Structure Reveals an Altered Intramolecular Hydrogen-bonding Pattern. Journal of Molecular Biology. 2005;351: 573–597. doi:10.1016/j.jmb.2005.05.053

9. Grasso V, Palermo S, Sierotzki H, Garibaldi A, Gisi U. Cytochrome b gene structure and consequences for resistance to Qo inhibitor fungicides in plant pathogens. Pest Management Science. 2006;62: 465–472. doi:10.1002/ps.1236

10. Fisher N, Meunier B. Molecular basis of resistance to cytochrome bc1 inhibitors. FEMS Yeast Research. 2008;8: 183–192. doi:10.1111/j.1567-1364.2007.00328.x

11. Dufour M-C, Fontaine S, Montarry J, Corio-Costet M-F. Assessment of fungicide resistance and pathogen diversity in Erysiphe necator using quantitative real-time PCR assays. Pest Management Science. 2011;67: 60–69. doi:10.1002/ps.2032

12. Miles LA, Miles TD, Kirk WW, Schilder AMC. Strobilurin (QoI) Resistance in Populations of Erysiphe necator on Grapes in Michigan. Plant Disease. 2012;96: 1621–1628. doi:10.1094/PDIS-01-12-0041-RE

13. Chen W-J, Delmotte F, Cervera SR, Douence L, Greif C, Corio-Costet M-F. At Least Two Origins of Fungicide Resistance in Grapevine Downy Mildew Populations. Applied and Environmental Microbiology. 2007;73: 5162–5172. doi:10.1128/AEM.00507-07

14. Ma Z, Felts D, Michailides TJ. Resistance to azoxystrobin in Alternaria isolates from pistachio in California. Pesticide Biochemistry and Physiology. 2003;77: 66–74. doi:10.1016/j.pestbp.2003.08.002

15. Pasche JS, Wharam CM, Gudmestad NC. Shift in Sensitivity of Alternaria solani in Response to QoI Fungicides. Plant Disease. 2004;88: 181–187. doi:10.1094/PDIS.2004.88.2.181

16. Amand O, Calay F, Coquillart L, Legat T, Bodson B, Moreau JM, et al. First detection of resistance to QoI fungicides in Mycosphaerella graminicola on winter wheat in Belgium. Commun Agric Appl Biol Sci. 2003;68: 519–531.

17. Torriani SF, Brunner PC, McDonald BA, Sierotzki H. QoI resistance emerged independently at least 4 times in European populations of Mycosphaerella graminicola. Pest Management Science. 2009;65: 155–162. doi:10.1002/ps.1662

18. Corio-Costet M-F. Fungicide resistance in Plasmopara viticola in France and anti resistance measures. Fungicide Resistance in crop protection : Risk and management. CABI Publishing; 2011. p. 304 p. Available: https://hal.inrae.fr/hal-02805838

19. Gisi U, Sierotzki H, Cook A, McCaffery A. Mechanisms influencing the evolution of resistance to Qo inhibitor fungicides. Pest Manag Sci. 2002;58: 859–867. doi:10.1002/ps.565

20. Cherrad S, Hernandez C, Steva H, Vacher S. Resistance of Plasmopara viticola to complex III inhibitors: a point on the phenotypic and genotypic characterization of strains. 12e Conférence Internationale sur les Maladies des Plantes, 11 et 12 décembre 2018, Tours, France. 2018; 449–459.

21. Fontaine S, Remuson F, Caddoux L, Barrès B. Investigation of the sensitivity of Plasmopara viticola to amisulbrom and ametoctradin in French vineyards using bioassays and molecular tools. Pest Management Science. 2019;75: 2115–2123. doi:10.1002/ps.5461

22. Fisher N, Meunier B, Biagini GA. The cytochrome bc1 complex as an antipathogenic target. FEBS Letters. 2020;594: 2935–2952. doi:10.1002/1873-3468.13868

23. Samils B, Andersson B, Edin E, Elfstrand M, Rönneburg T, Bucur D, et al. Development of a PacBio Long-Read Sequencing Assay for High Throughput Detection of Fungicide Resistance in Zymoseptoria tritici. Frontiers in Microbiology. 2021;12. Available: https://www.frontiersin.org/article/10.3389/fmicb.2021.692845

24. Pereira D, McDonald BA, Croll D. The Genetic Architecture of Emerging Fungicide Resistance in Populations of a Global Wheat Pathogen. Genome Biol Evol. 2020;12: 2231– 2244. doi:10.1093/gbe/evaa203

25. Waterhouse AM, Procter JB, Martin DMA, Clamp M, Barton GJ. Jalview Version 2— a multiple sequence alignment editor and analysis workbench. Bioinformatics. 2009;25: 1189– 1191. doi:10.1093/bioinformatics/btp033

26. Li H. Minimap2: pairwise alignment for nucleotide sequences. Bioinformatics. 2018;34: 3094–3100. doi:10.1093/bioinformatics/bty191

27. Garrison E, Marth G. Haplotype-based variant detection from short-read sequencing. arXiv:12073907 [q-bio]. 2012 [cited 28 Mar 2022]. Available: http://arxiv.org/abs/1207.3907

28. Garrison E, Kronenberg ZN, Dawson ET, Pedersen BS, Prins P. Vcflib and tools for processing the VCF variant call format. bioRxiv; 2021. p. 2021.05.21.445151. doi:10.1101/2021.05.21.445151

29. Camacho C, Coulouris G, Avagyan V, Ma N, Papadopoulos J, Bealer K, et al. BLAST+: architecture and applications. BMC Bioinformatics. 2009;10: 421. doi:10.1186/1471-2105-10-421

30. Fehr M, Wolf A, Stammler G. Binding of the respiratory chain inhibitor ametoctradin to the mitochondrial bc1 complex. Pest Management Science. 2016;72: 591–602. doi:10.1002/ps.4031

31. Dreinert A, Wolf A, Mentzel T, Meunier B, Fehr M. The cytochrome bc1 complex inhibitor Ametoctradin has an unusual binding mode. Biochimica et Biophysica Acta (BBA) -Bioenergetics. 2018;1859: 567–576. doi:10.1016/j.bbabio.2018.04.008

32. Lucas JA, Hawkins NJ, Fraaije BA. Chapter Two - The Evolution of Fungicide Resistance. In: Sariaslani S, Gadd GM, editors. Advances in Applied Microbiology. Academic Press; 2015. pp. 29–92. doi:10.1016/bs.aambs.2014.09.001

33. Mounkoro P, Michel T, Benhachemi R, Surpateanu G, Iorga BI, Fisher N, et al. Mitochondrial complex III Qi-site inhibitor resistance mutations found in laboratory selected mutants and field isolates. Pest Management Science. 2019;75: 2107–2114. doi:10.1002/ps.5264

34. Toffolatti SL, Serrati L, Sierotzki H, Gisi U, Vercesi A. Assessment of QoI resistance in Plasmopara viticola oospores. Pest Management Science. 2007;63: 194–201. doi:10.1002/ps.1327

35. Malandrakis AA, Markoglou AN, Nikou DC, Vontas JG, Ziogas BN. Molecular diagnostic for detecting the cytochrome b G143S – QoI resistance mutation in Cercospora beticola. Pesticide Biochemistry and Physiology. 2011;100: 87–92. doi:10.1016/j.pestbp.2011.02.011

36. Martinelli F, Scalenghe R, Davino S, Panno S, Scuderi G, Ruisi P, et al. Advanced methods of plant disease detection. A review. Agron Sustain Dev. 2015;35: 1–25. doi:10.1007/s13593-014-0246-1

37. Rallos LEE, Baudoin AB. Co-Occurrence of Two Allelic Variants of CYP51 in Erysiphe necator and Their Correlation with Over-Expression for DMI Resistance. PLOS ONE. 2016;11: e0148025. doi:10.1371/journal.pone.0148025

38. Massi F, Torriani SFF, Borghi L, Toffolatti SL. Fungicide Resistance Evolution and Detection in Plant Pathogens: Plasmopara viticola as a Case Study. Microorganisms. 2021;9: 119. doi:10.3390/microorganisms9010119

39. Blum M, Waldner M, Gisi U. A single point mutation in the novel PvCesA3 gene confers resistance to the carboxylic acid amide fungicide mandipropamid in Plasmopara viticola. Fungal Genetics and Biology. 2010;47: 499–510. doi:10.1016/j.fgb.2010.02.009

40. Santos RF, Fraaije BA, Garrido L da R, Monteiro-Vitorello CB, Amorim L. Multiple resistance of Plasmopara viticola to QoI and CAA fungicides in Brazil. Plant Pathology. 2020;69: 1708–1720. doi:10.1111/ppa.13254

41. Rehfus A, Miessner S, Achenbach J, Strobel D, Bryson R, Stammler G. Emergence of succinate dehydrogenase inhibitor resistance of Pyrenophora teres in Europe. Pest Management Science. 2016;72: 1977–1988. doi:10.1002/ps.4244

42. Zhou T, Mehl HL. Rapid quantification of the G143A mutation conferring fungicide resistance in Virginia populations of Cercospora sojina using pyrosequencing. Crop Protection. 2020;127: 104942. doi:10.1016/j.cropro.2019.104942

43. Assinger T, Fountaine J, Torriani S, Accardo S, Bernhard-Frey R, Gottula J, et al. Detection of Ramularia collo-cygni DMI- and SDHI-resistant field populations in Austria and the effect of fungicides on the population and genetic diversity. Eur J Plant Pathol. 2022;162: 575–594. doi:10.1007/s10658-021-02422-5

44. Hendriksen RS, Bortolaia V, Tate H, Tyson GH, Aarestrup FM, McDermott PF. Using Genomics to Track Global Antimicrobial Resistance. Frontiers in Public Health. 2019;7. Available: https://www.frontiersin.org/article/10.3389/fpubh.2019.00242

45. Berbers B, Saltykova A, Garcia-Graells C, Philipp P, Arella F, Marchal K, et al. Combining short and long read sequencing to characterize antimicrobial resistance genes on plasmids applied to an unauthorized genetically modified Bacillus. Sci Rep. 2020;10: 4310. doi:10.1038/s41598-020-61158-0

46. Taxt AM, Avershina E, Frye SA, Naseer U, Ahmad R. Rapid identification of pathogens, antibiotic resistance genes and plasmids in blood cultures by nanopore sequencing. Sci Rep. 2020;10: 7622. doi:10.1038/s41598-020-64616-x

47. Liefting LW, Waite DW, Thompson JR. Application of Oxford Nanopore Technology to Plant Virus Detection. Viruses. 2021;13: 1424. doi:10.3390/v13081424

48. Kankanala P, Nandety RS, Mysore KS. Genomics of Plant Disease Resistance in Legumes. Frontiers in Plant Science. 2019;10. Available: https://www.frontiersin.org/article/10.3389/fpls.2019.01345

49. Pieczul K, Wąsowska A. The application of next-generation sequencing (NGS) for monitoring of Zymoseptoria tritici QoI resistance. Crop Protection. 2017;92: 143–147. doi:10.1016/j.cropro.2016.10.026

50. Van Leeuwen T, Vanholme B, Van Pottelberge S, Van Nieuwenhuyse P, Nauen R, Tirry L, et al. Mitochondrial heteroplasmy and the evolution of insecticide resistance: Non-Mendelian inheritance in action. Proceedings of the National Academy of Sciences. 2008;105: 5980–5985. doi:10.1073/pnas.0802224105

51. Hu M, Chen S. Non-Target Site Mechanisms of Fungicide Resistance in Crop Pathogens: A Review. Microorganisms. 2021;9: 502. doi:10.3390/microorganisms9030502

52. Mosquera S, Chen L-H, Aegerter B, Miyao E, Salvucci A, Chang T-C, et al. Cloning of the Cytochrome b Gene From the Tomato Powdery Mildew Fungus Leveillula taurica Reveals High Levels of Allelic Variation and Heteroplasmy for the G143A Mutation. Frontiers in Microbiology. 2019;10. Available: https://www.frontiersin.org/article/10.3389/fmicb.2019.00663

53. Magnien C, Micoud A, Glain M, Remuson F. QoI resistance of downy mildew-monitoring and tests 2002. Ann 7th Int Conf Plant Dis, Tours, France(On CD-ROM). 2003.

54. Salipante SJ, Kawashima T, Rosenthal C, Hoogestraat DR, Cummings LA, Sengupta DJ, et al. Performance Comparison of Illumina and Ion Torrent Next-Generation Sequencing Platforms for 16S rRNA-Based Bacterial Community Profiling. Applied and Environmental Microbiology. 2014;80: 7583–7591. doi:10.1128/AEM.02206-14

55. Boykin L, Ghalab A, Marchi BRD, Savill A, Wainaina JM, Kinene T, et al. Real time portable genome sequencing for global food security. F1000Research; 2018. doi:10.12688/f1000research.15507.1

56. Boykin LM, Sseruwagi P, Alicai T, Ateka E, Mohammed IU, Stanton J-AL, et al. Tree Lab: Portable Genomics for Early Detection of Plant Viruses and Pests in Sub-Saharan Africa. Genes. 2019;10: 632. doi:10.3390/genes10090632

57. Shaffer L. Portable DNA sequencer helps farmers stymie devastating viruses. Proceedings of the National Academy of Sciences. 2019;116: 3351–3353. doi:10.1073/pnas.1901806116

58. Salcedo AF, Purayannur S, Standish JR, Miles T, Thiessen L, Quesada-Ocampo LM. Fantastic Downy Mildew Pathogens and How to Find Them: Advances in Detection and Diagnostics. Plants. 2021;10: 435. doi:10.3390/plants10030435

